# The ascending arousal system shapes low-dimensional neural dynamics to mediate awareness of intrinsic cognitive states

**DOI:** 10.1101/2021.03.30.437635

**Authors:** Brandon Munn, Eli J. Müller, Gabriel Wainstein, James M. Shine

**Affiliations:** Complex Systems Research Group, The University of Sydney, Sydney, NSW, Australia; Brain and Mind Centre, The University of Sydney, Sydney, NSW, Australia

## Abstract

Models of cognitive function typically focus on the cerebral cortex and hence overlook functional links to subcortical structures. This view neglects the highly-conserved ascending arousal system’s role and the computational capacities it provides the brain. In this study, we test the hypothesis that the ascending arousal system modulates cortical neural gain to alter the low-dimensional energy landscape of cortical dynamics. Our analyses of spontaneous functional magnetic resonance imaging data and phasic bursts in both locus coeruleus and basal forebrain demonstrate precise time-locked relationships between brainstem activity, low-dimensional energy landscapes, network topology, and spatiotemporal travelling waves. We extend our analysis to a cohort of experienced meditators and demonstrate locus coeruleus-mediated network dynamics were associated with internal shifts in conscious awareness. Together, these results present a novel view of brain organization that highlights the ascending arousal system’s role in shaping both the dynamics of the cerebral cortex and conscious awareness.

## Main Text

It is often difficult to see the forest for the trees, but to fully understand a concept typically involves an accurate depiction of both. That is, we need to comprehend not only the detailed workings of a specific system, but also how that system functions within a broader context of interacting parts. Modern theories of whole-brain function exemplify this challenge. For instance, activity in the brain has been shown to incorporate signatures of both local computational specificity (e.g., specialized regions within the cerebral cortex) as well as system-wide integration (e.g., the interactions between the cortex and the rest of the brain)^1,2^. Anatomical evidence suggests that the balance between integration and segregation is mediated in part by the relatively fixed white matter connections between cerebral cortical regions^1^ — local connectivity motifs support segregated activity, whereas the axonal, reentrant connections between regions act to integrate the distributed signals via a highly interconnected structural backbone^3^. However, how the human brain is also capable of remarkable contextual flexibility given this relatively fixed connectivity remains poorly understood.

During cognitive tasks, neural activity rapidly reconfigures the functional large-scale network architecture of the brain to facilitate coordination between otherwise segregated cortical regions. Precisely how this flexibility is implemented in the brain without altering structural connectivity remains an open question in systems neuroscience. Although it is often overlooked in theories of whole brain function, the neuromodulatory ascending arousal system is well-placed to mediate this role^4^. The arousal system is comprised of a range of nuclei spread across the brainstem and forebrain that send wide-reaching axons to the rest of the central nervous system^5^. At their target sites, arousal neurons release neuromodulatory neurotransmitters that shape and constrain a region’s processing mode — altering their excitability and responsivity without necessarily causing them to fire an action potential^4,6^. As a result, subtle changes in the concentration of neuromodulatory chemicals can cause massive alterations in the dynamics of the target regions, leading to nonlinear effects on the coordinated patterns of activity that emerge from ‘simple’ neuronal circuits^4^.

The ascending arousal system also contains substantial heterogeneity — unique cell populations project in diverse ways to the cerebral cortex and release distinct neurotransmitters. One key dichotomy is the distinction between adrenergic neuromodulation (predominantly via the locus coeruleus [LC]), which promotes arousal and exploratory behaviour^7^, and cholinergic neuromodulation (such as via the basal nucleus of Meynert [BNM]), which is associated with attentional focus and vigilance^8^. These highly interconnected^9^ structures both promote wakefulness and arousal^10,11^, albeit via distinct topological projections to the cerebral cortex: the LC projects in a diffuse manner that crosses typical specialist boundaries, whereas the BNM projects in a more targeted, regionspecific manner^12^ (Fig. 1A). The two systems have also been linked with distinct and complimentary computational principles: the noradrenergic LC is presumed to modulate interactions between neurons (response gain)^13^, whereas the cholinergic BNM is presumed to facilitate divisive normalization (multiplicative gain)^14^. Based on these anatomical and computational features, we have hypothesized that the interaction between these two neuromodulatory systems is crucial for mediating the dynamic, flexible balance between integration and segregation in the brain^15^.

**Figure 1.**
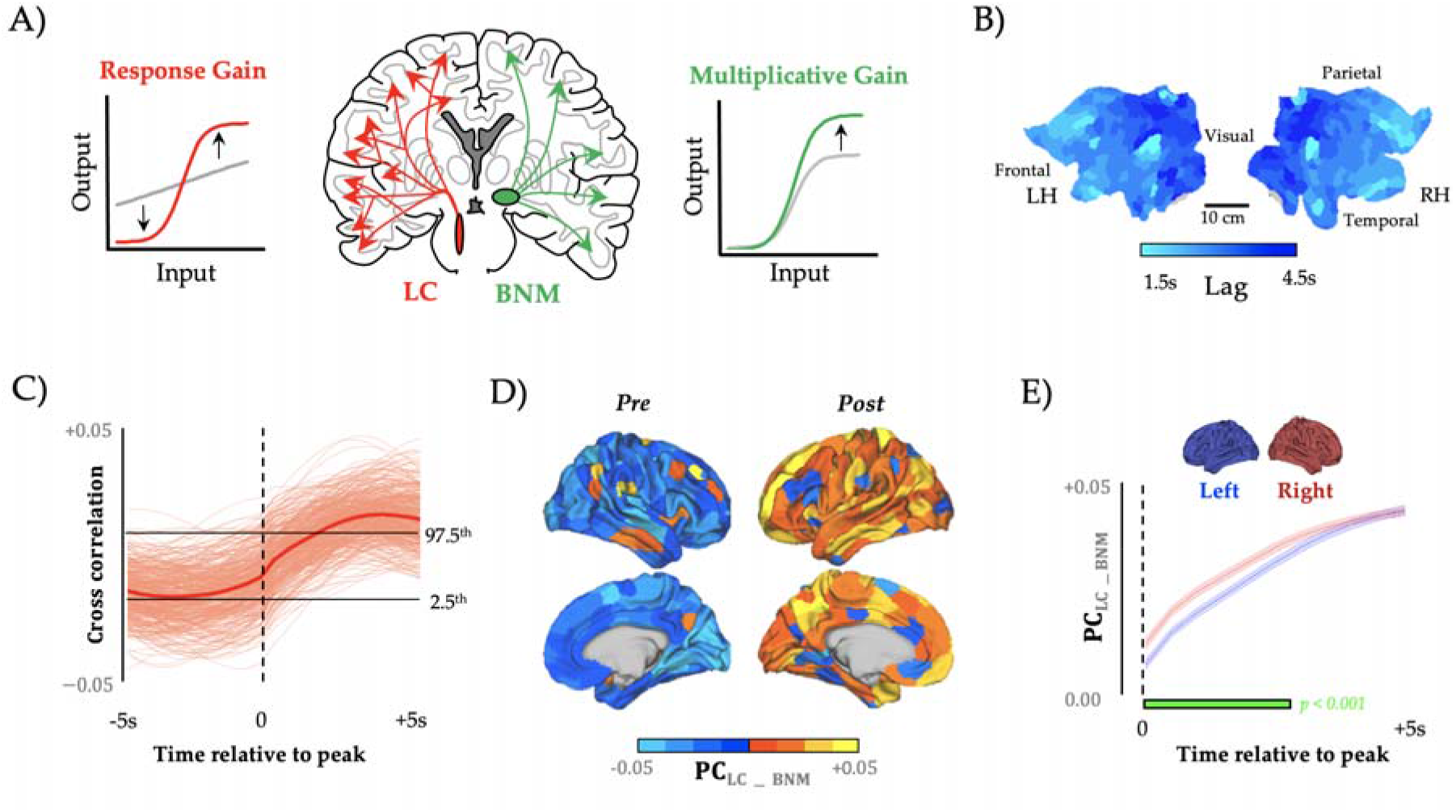
Sympathetic activity precedes network-level integration. A) regional time series were extracted from the locus coeruleus (red), which is thought to alter response gain, and the basal nucleus of Meynert (green), which is thought to alter multiplicative gain, and compared to BOLD signal and topological signatures during the resting state; B) we observed a anterior-to-posterior traveling wave (velocity ~ 0.13ms^-1^) following peaks in τ_LC-BNM_ which are shown on both the left (LH) and right (RH) hemispheres of a cortical flat map; C) the lagged cross-correlation between and PC — dotted line depicts the zero-lag correlation, and the black lines depict the upper (lower) bounds of a block-resampled null model (95% CI); D) mean cortical participation coefficient (PC) preceding (left) and following (right) the zero-lagged value, only the right hemisphere is shown (mirrored for ‘Post’); E) the participation coefficient following peak was higher in the right- (red) vs. the left- (blue) hemisphere (p < 0.001; green bar).

Another crucial feature of the ascending arousal system is that the number of neurons that project to the cerebral cortex is several orders of magnitude smaller than those that project back to the brainstem and forebrain^16–18^. Based on this feature, we further hypothesize that shifts in arousal are realized through a low-dimensional modulation of the ongoing neural activity (‘brain state’)^17^. Conceptually, low-dimensional neural dynamics can be depicted as evolving on a brain state energy landscape^19^, where the energy of a given state corresponds to the occurrence probability, e.g. high energy brain states have a low occurrence probability (and *v. v*). That is brain states evolve along the energy landscape topography, much like a ball rolls under the influence of gravity down a valley and requires energy to traverse up a hill, this corresponds to an evolution towards an attractive or repulsive brain state, respectively. This technique can resolve what might otherwise be obscured states of attraction (and repulsion) in a multi-stable system and has been successfully applied to the dynamics of spiking neurons^20,21^, blood oxygenation level dependent (BOLD) functional magnetic resonance imaging (fMRI)^22,23^, and magnetoencephalography (MEG)^24^. The approach offers several conceptual advances, but perhaps most importantly, it renders the otherwise daunting task of systems-level interpretation relatively intuitive. Importantly, this framework is not a mere analogy^30^, as the topography of the energy landscape shares a 1-to-1 correspondence with the generative equations required to synthesize realistic neural timeseries data^25^. In this manuscript, we test these ideas by combining high-resolution resting state fMRI data with analytic techniques from the study of complex systems.

## Results

To begin with, we extracted time series data from major subcortical hubs within the noradrenergic LC^9^ (Fig. 1A, red) and cholinergic BNM^26^ (Fig. 1A, green) systems from 59 healthy participants who had undergone high-resolution, 7T resting-state functional magnetic resonance imaging (fMRI; 2 mm^3^ voxels; TR = 586 ms repetition time). Given the known spatiotemporal interactions between the ascending arousal system and fluctuations in cerebrospinal fluid, we first controlled for activity fluctuations in the nearby fourth ventricle, which contains no neural structures, but nonetheless can cause alterations in the BOLD signal over time. We next accounted for nearby gray-matter signals, by regressing the signal from the nearby pontine nuclei. Using the residuals from these regressions from the LC signal, τ_LC_, and the BMN signal, τ_BNM_, we focused on the difference between these signals (τ_LC-BNM_ and τ_BNM-LC_ concatenated across subjects) and then identified time points associated with phasic bursts of LC activity that led to sustained adrenergic (versus cholinergic) influence over evolving brain state dynamics (and *v.v.* for phasic bursts of BNM; see Methods). Importantly, the phasic mode of firing within the noradrenergic arousal system has been specifically linked to systemic influences that occur on time-scales relevant to cognitive function^8,27^. Tracking the mean cortical BOLD response around these peaks identified a spatiotemporal travelling wave (Fig. 1B; velocity = 0.13ms^-1^) that propagated from frontal to sensory cortices and tracked closely with the known path of the dorsal noradrenergic bundle^9^, albeit with a preserved ‘island’ within the parietal operculum (Fig. 1B). A block-resampling null model was applied to ensure that the results were not due to spatial-autocorrelation (p < 0.05; see Methods). These results can be inverted for BNM activity (relative to LC) as τ_BNM-LC_ = -τ_LC-BNMO_ Furthermore, these results confirm that coordinated macroscale activity patterns align to fluctuations in activity within the ascending arousal system of the brainstem^28^.

### Time-varying network topology

Based on previous empirical^29^, modelling^30^ and theoretical^15^ work, we predicted that phasic bursts in τ_LC-BNM_ would facilitate network-level integration by modulating increased neural gain among regions distributed across the cerebral cortex. As predicted, we observed a strong positive correlation between τ_LC-BNM_ and network-level integration (p < 0.05, block-resampling null model; Fig. 1C) across the brain (Fig. 1D). An increase in phasic activity within the LC (relative to the BNM) preceded an increase in the mean level of integration within the cerebral cortex that was dominated by the frontoparietal cortices (Fig. 1D; parcellated according to the 17 resting-state networks identified in^31^). Interestingly, this global integration was opposed by a relative topological segregation of limbic, visual, and motor cortices (Fig. S2). This increase in the synchronisation of the frontoparietal cortices following an increase in sensory-limbic coordination and LC activity may reflect arousal- enhanced processing of sensory stimuli^32,33^. Furthermore, regional integration occurred earlier in the right- vs. the left-hemisphere (p<0.001; Fig. 1E), which is consistent with the known anatomical bias of the LC system^34,35^. Together, these findings provide robust evidence for the hypothesis that the balance between ascending noradrenergic and cholinergic tone facilitates a transition towards topological integration across the frontoparietal network of the brain^15^.

### Neuromodulation of the Energy Landscape

The results of our initial analysis demonstrate that coordinated distributed activity in the cortex align with changes in small groups of neuromodulatory cells activity in the brainstem and forebrain, which in turn are proposed to constrain brain dynamics onto a low-dimensional energy landscape (Fig. 2A). The effects of noradrenaline and acetylcholine can also be easily viewed through this lens: by integrating the brain, noradrenaline should flatten the energy landscape (Fig. 2A, red) facilitating otherwise unlikely brain state transitions, whereas in contrast, the segregative nature of cholinergic activity should act to deepen energy valleys (Fig. 2A, green) decreasing the likelihood of a brain state transition. In previous work, we have shown a correspondence between low-dimensional brain state dynamics across multiple cognitive tasks and the heterogenous expression of metabotropic neuromodulatory receptors^17^. This implies that neuromodulators act similar to catalysts in chemical reactions, which lower (or raise) the activation energy (E_A_) required to transform chemicals from one steady state (or energy well) to another (Fig. 2B). In the context of the interconnected, heterarchical networks that comprise the cerebral cortex, this would have the effect of flattening (or deepening) the energy landscape, promoting variable (or rigid) brain states^36^ (Fig. 2A).

**Figure 2.**
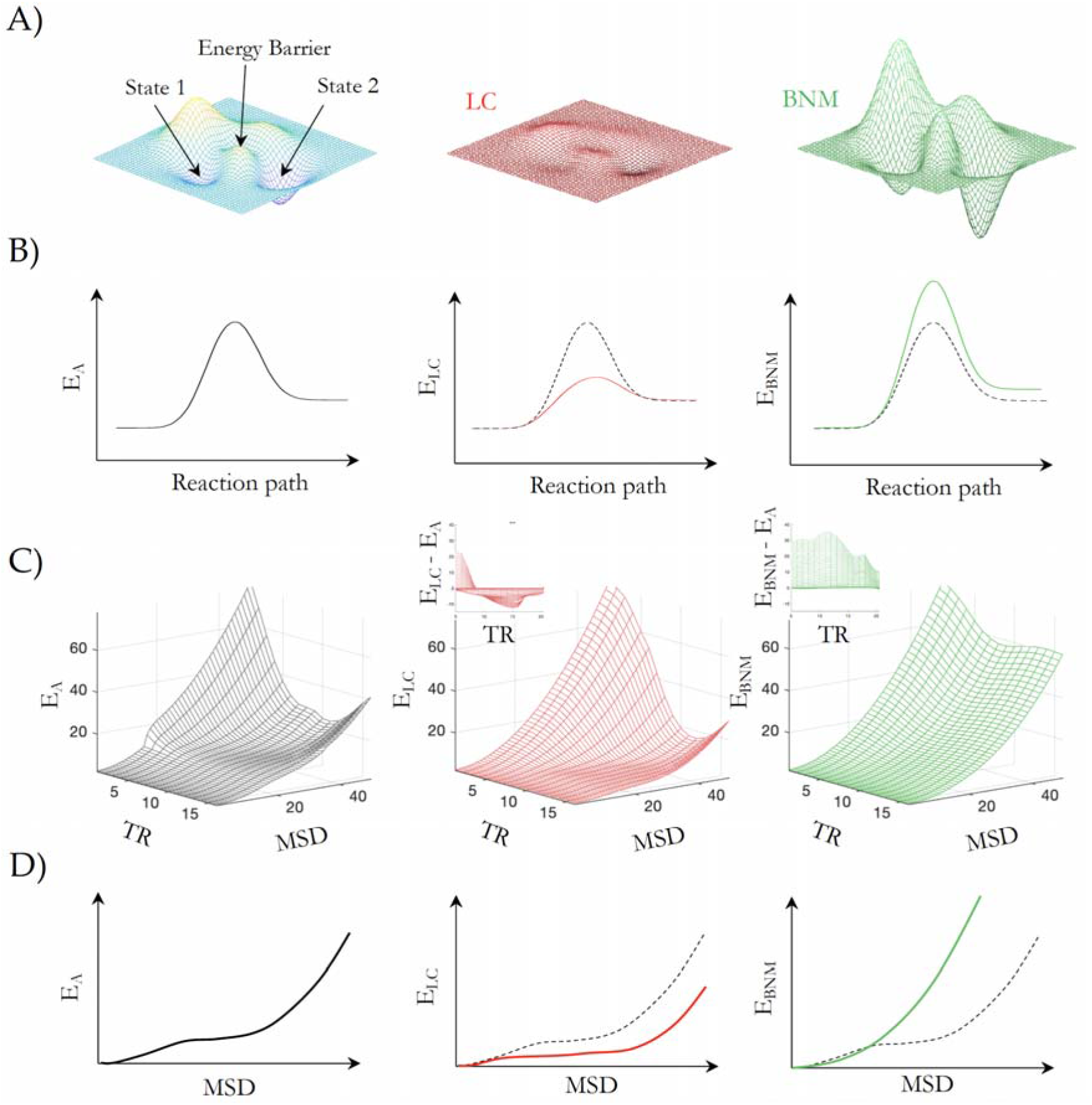
LC and BNM mediated shifts in energy landscape brain state space dynamics. A) an energy landscape, which defines the energy required to move between different brain states: by increasing response gain, the LC should flatten the energy landscape (red); by increasing multiplicative gain, the BNM should deepen the energy wells (green); B) the topography of the energy landscape can be conceptualized as similar to the activation energy (E_A_) that must be overcome in order to convert one chemical to another; C) Empirical BOLD trajectory energy as a function of MSD and TR of the baseline activity (E_A_, black) and after phasic bursts in LC (E_LC_, red) and BNM (E_BNM_, green) — relative to the baseline energy landscape phasic bursts in LC (E_LC_ - E_A_, red inset) lead to a flattening or reduction of the energy landscape, whereas peaks in BNM (E_BNM_ - E_A_, green inset) lead to a raising of the energy landscape. D) Empirical activation energy as a function of MSD averaged over lags TR during base baseline activity (E_A_, Left) and following phasic bursts in LC (E_LC_, red) and BNM (E_BNM_, green).

To elucidate the role of phasic activity from the neuromodulatory system in modifying the energy landscape, we first estimated the energy of BOLD signal transitions across the cerebral cortex. Importantly, the term ‘energy’ here is used in reference to its definition in statistical physics and hence does not represent the biological use of the term, which instead stands for the energy used by the brain to maintain or change neural activity. Specifically, we define the energy landscape, ***E***, as the natural logarithm of the inverse probability of observing a given BOLD ***MSD*** at a given time-lag ***t, P***(***MSD, t***), calculated as 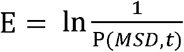 where 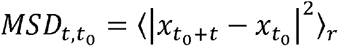 is the mean-squared displacement (MSD) of BOLD signal, ***x_t_*** = [***x*_1,*t*_,*x*_2,*t*_, …, *x_r,t_***] across ***r*** voxels and ***t*** is the number of time-lags of size TR from a reference timepoint ***t*_0_**^20^. The probability of a BOLD signal transition, ***P***(***MSD, t***), was estimated from the sampled *MSD_t to_,* and we used a Gaussian kernel density estimation 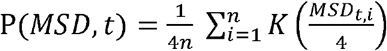, where 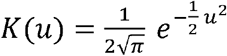 (see Methods). Our analysis is consistent with the statistical mechanics interpretation that the energy of a given state, ***E_σ_***, and its probability are related 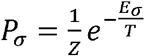, where ***Z*** is the normalisation function and ***T*** is a scaling factor equivalent to temperature in thermodynamics, where we set ***T* = 1** and ***Z* = 1**^20^. In this framework, a highly probable relative change in BOLD (as quantified by the MSD) corresponds to a relatively low energy transition (i.e., low E_*A*_), whereas an infrequently visited state will require the most energy (i.e., high E_*A*_).

By treating energy as inversely proportional to the probability of brain state occurrence, our approach resembles other studies that have been applied to spiking dynamics of neuronal populations, spiking neurons^20,21^, BOLD fMRI^22,23^, MEG^24^, and natural scene^37^. However, these studies binarized continuous signals to reduce the brain state space (to 2^r^ states), however this approach requires the fitting of a threshold, which can be problematic in continuously recorded data. In contrast, our approach reduces the dimensionality by analysing the likelihood of a change in BOLD activity (i.e., the MSD), and thus retains the dimensionality of the underlying signal without the need for thresholding. This approach overcomes a major limitation inherent to previous energy landscape studies that require a large sample size to sufficiently sample the brain state space.

With this in mind, we turned our attention to the relationship between the ascending arousal system dynamics and the MSD energy landscape. To test the hypothesis that the neuromodulatory system alters the topography of the energy landscape, we calculated BOLD MSD energetics following phasic bursts of both LC relative to BNM (***τ_LC-BNM_***), ***E_LC_*** and BNM relative to LC (*τ_BNM-LC_*), ***E_BNM_***, i.e., ***t*_0_** was the onset of a phasic burst, and compared these to sampled brain evolutions without large changes in LC and BNM arousal i.e., ***t*_0_** was all timepoints outside of a phasic burst in LC and BNM, analogous to the baseline energy landscape ***E_A_***. We identified phasic bursts as peaks in the second derivative of the arousal BOLD signals ***τ_LC-BNM_*** and ***τ_BNM-LC_*** that lead to a sustained increase in BOLD activity for each individual (see Methods) and using these criteria, we identified 148 τ_LC-BNM_ time points and 130 τ_BNM-LC_ time points.

The energy landscapes for these three states are defined by the energy for a given BOLD MSD at a given TR delay. Figure 2C demonstrates the baseline energy landscape (Fig. 2C, black), which corresponds to the reaction pathway in Fig. 2B, and the MSD energy landscape following phasic bursts in the LC (Fig. 2C, red) and BNM (Fig. 2C, green). These figures demonstrate an MSD energy landscape across displacement and time, wherein the energy relates to the likelihood of seeing a given mean change in bold activity (i.e., MSD) at a given temporal displacement (i.e., TR). For example, all the MSD energy landscapes have a high energy peak for large MSD at a short timescale as it is extremely unlikely that the bold activity would change significantly (quantified by a large MSD) in one TR (~0.5s), and neuromodulation increases the energy of such an initial change. The utility of the MSD energy landscape can be seen when comparing large phasic bursts of LC and BNM relative to baseline fluctuations. We found the largest change occurs around 10-15 TR (~6-9s) following a phasic burst that typically corresponds to a peak in the LC or BNM BOLD signal. At this ~6-9s temporal delay we see direct evidence that a phasic burst of LC flattened the energy landscape (decreased the energy relative to baseline Fig. 2C, red inset), thus making previously unlikely large MSD trajectories far more probable (Fig. 2D red), whereas a phasic burst of BNM activity (increased energy relative to baseline Fig. 2C, green inset) caused the energy landscape to be elevated, thus promoting local trajectories, and making large MSD deviations unlikely (Fig. 2D green). These patterns are analogous to modulating a physical landscape in which towns sit within valleys separated by impassable mountains — when BNM is high, the towns remain isolated, whereas when LC is high, the towns are separated by easily navigated rolling plains and transitions between towns (novel combinations of consecutive brain states) can be easily realised.

We next asked whether LC and BNM combined synergistically to alter the energy landscape. To achieve this, we isolated simultaneous phasic peaks in both LC and BNM (*τ_LC-BNM_*) We found that the LC + BNM energy landscape differed from either independent LC or BNM activation, shifting the brain state into divergent regimes than could be explained by the HRF. By comparing the MSD energy topography for a given TR slice we found that the landscape switched from an anti- to de-correlation with the HRF. In other words, the cooperative behaviour between the noradrenergic and cholinergic systems allowed the brain to reach unique BOLD MSDs that neither could facilitate individually. To examine how simultaneous LC+BNM activity altered the energy landscape, we compared the energy relative to the two individual landscapes. As demonstrated in Fig. S3, the energy landscape following phasic bursts of LC+BNM differed in magnitude from that expected from a linear superposition of the LC and BNM energy landscape — i.e., LC+BNM ≠ (LC) + (BNM). Furthermore, to explore the dominance of either LC or BNM in this signal, we minimised the relationship LC+BNM = *α*LC + ***β***BNM (conditional upon ***α*** and ***β*** being positive constants) and found that ***α* = 0.16** and ***β* = 0.84** gave the best match to the LC+BNM energy landscape. That is, the BNM dynamics dominates the simultaneous LC+BNM energy landscape, which is consistent with the unidirectional synaptic projections from the LC that synapse upon the BNM on their way through to the cortex^12^, and suggests that phasic LC+BNM bursts may be initiated by the LC in order to elicit a cascade of BNM activity.

### Conscious awareness of shifts in BOLD state

Interpreting the relationship between neuroimaging data and conscious awareness is notoriously challenging. For instance, it is currently not possible to directly determine the contents of self-directed thought without intervening, and thus altering, the contents of consciousness^38^. Although we can’t determine the contents of consciousness directly, we can use task designs to modulate the state of consciousness. To this end, we leveraged data from a group of 14 expert meditators who were asked to meditate during an fMRI scanning session^39^, and to press a button when they noticed that their focus had drifted from their breath (Fig. 3A). At this point, there is a mismatch between expectation and conscious awareness, which is an internal state that has been previously linked to the activation of the noradrenergic system, both in theoretical^40,41^ and computational^42^ work. Based on these studies, we predicted that the switch in internal conscious awareness would be facilitated by increases in locus coeruleus-mediated integration and subsequent reconfiguration of low-dimensional brain states. Analysing time-resolved network data with a finite impulse response model, we observed a peak in locus coeruleus activity (Fig. 3B), TR-to-TR mean squared displacement (Fig. 3C) and elevated network-level integration (Fig. 3D) surrounding the change in conscious awareness (all p_*PE*R*M*_ < 0.05; 95% CI of null distribution). These results confirm that the locus coeruleus mediates energy landscape reconfigurations and that these changes modulate internal states of conscious awareness.

**Figure 3.**
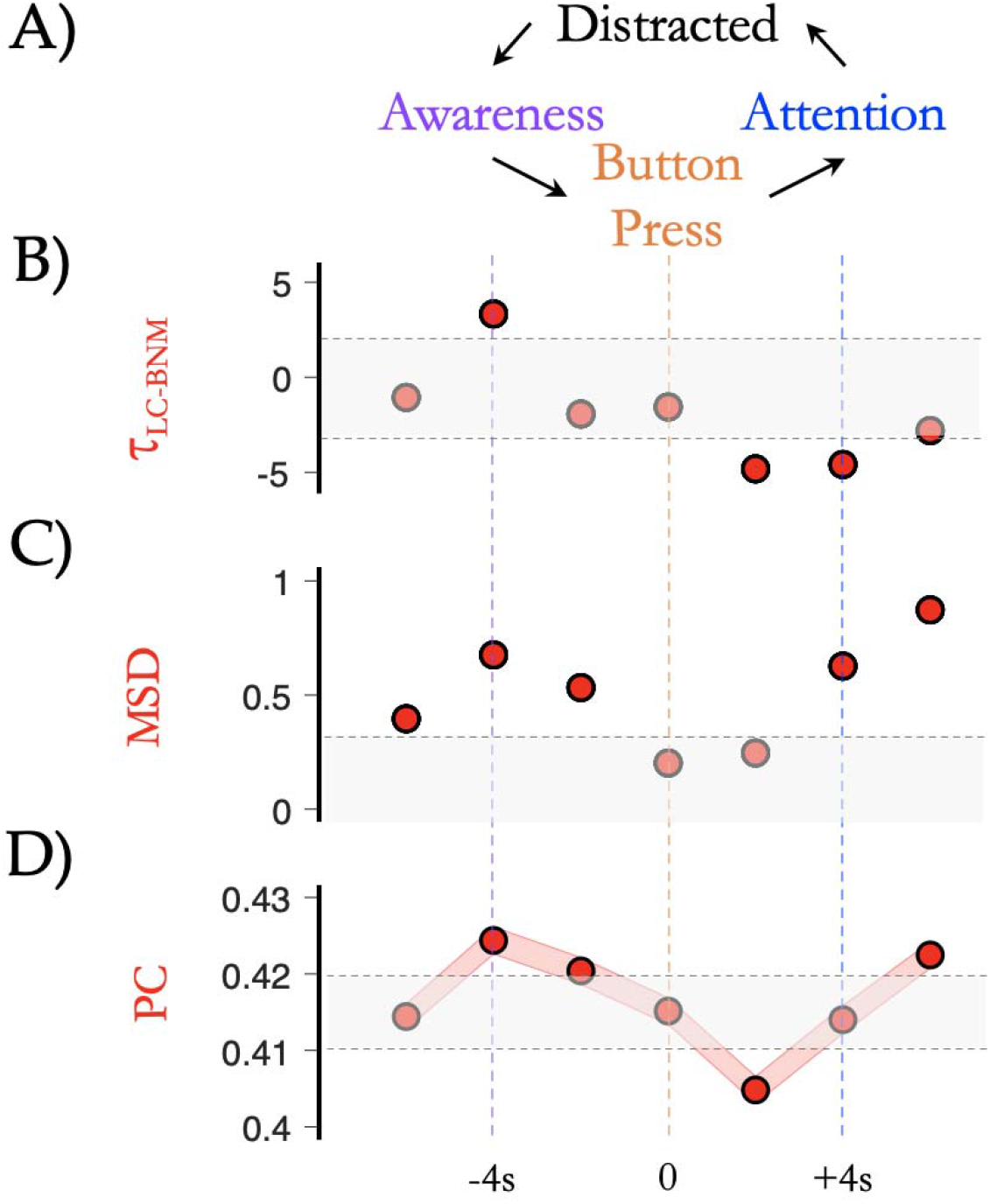
Awareness of intrinsic state changes. A) participants performing breath-awareness meditation (Focus; blue) were trained to respond with a Button Press (orange) when they became Aware (purple) that they had become Distracted (i.e., their attention had wandered from their breath) and to then re-focus their Attention (blue) on their breath; B) we observed a peak in ***τ**_LC-BNM_* ftc., LC > BNM; red) ~4 seconds before the button press, which then returned to low levels (i.e., BNM > LC) in the 2-4 seconds following the button press; C) the Mean Squared Displacement (MSD; dark orange) of TR-to-TR BOLD signal was increased above null values around the peak in ***τ**_LC-BNM_*, as well as following the re-establishment of attentional focus (in panels B & C, grey shading depicts 95% CI of block-resampled null distribution); D) we observed a peak in mean participation coefficient (PC) ~4 seconds (2 TRs) prior to the Button Press during the task. All: grey shading depicts 97.5^th^ and 2.5^th^ percentile of null distribution i.e., outside grey shading indicates a value different than null [p < 0.05]; and lower: red shading represents SEM error bars).

### Discussion

Our results provide evidence for an arousal-mediated macroscopic network and energy landscape reconfiguration which tracks with moment-to-moment alterations in conscious awareness. By tracking fluctuations in BOLD signal within the noradrenergic LC and the cholinergic BNM, we were able to demonstrate fundamental ways in which the low-dimensional, dynamic, and topological signature of cortical dynamics was related to changes within the ascending arousal system. Furthermore, we demonstrated a link between these dynamic reconfigurations and alterations in conscious awareness in a cohort of experienced meditators. In this way, our results provide a novel, systems-level perspective on the distributed dynamics of the human brain.

There is growing evidence that distributed neural dynamics in the brain are well described by relatively low-dimensional models^17,18,43–45^, however the biological constraints that impose these features on the brain remain poorly understood. Due to the low number of cells in the arousal system and their broad projections to the rest of the brain, we theorized that neuromodulatory regions are well-placed to shape and constrain the vast number of neurons in the cerebral cortex into low-dimensional dynamic modes. Our results support this prediction by showing that patterns of activity in key regions within the brainstem and forebrain relate to fundamental alterations in a dynamically evolving energy landscape. In other words, neural state space trajectories are a powerful framework that extends beyond that of mere analogy, and the ascending arousal system is well-placed to mediate deformations in the energy landscape.

Much in the same way that there are many different reference frames for navigation — e.g., egocentric (i.e., straight, left, right directions), which is independent of the environment; and allocentric (i.e., following compass directions and visual cues), which is dependent on the environment — we can interrogate energy landscapes using different vantage points on BOLD dynamics. Our displacement framework is consistent with an egocentric (or ‘first-person’) frame of reference, wherein MSD is used to track BOLD trajectories from an initial state which maps out the topology of the energy landscape (i.e., a BOLD MSD implies a BOLD trajectory). Nonetheless, the method does not distinguish between two different neural trajectories that possess the same MSD. In comparison, other methods have evaluated the energy landscape for a given pre-defined state estimated from thresholded BOLD timeseries^22,23^ a framework consistent with an allocentric (or ‘third-person’) reference frame. This framework has the advantage of calculating energy for a given state, however, it also requires substantial exploration of the state-space — which is typically unfeasible — or the need to resort to severe coarse-graining (such as the binarization of BOLD activity) which further diminishes interpretability. Furthermore, the allocentric view does not provide insights into the transitions between each energy state, whereas this information is inherent to the egocentric reference. Along these lines, we found that the egocentric reference frame clearly demonstrated the flattening and deepening of the energy landscape, providing indirect evidence that the ascending arousal system is well set-up to control brain-state dynamics ‘egocentrically’ (as opposed to specific neural activity patterns). Nevertheless, given improvements in recording length and novel analytic techniques to probe the brains dynamical landscape, we expect that the field will ultimately discover even more optimal mappings between neurobiology and low-dimensional brain state dynamics.

The results of our state-space analysis have important implications for the biological mechanisms underlying cognition. For instance, the concept of locus coeruleus-mediated energy landscape flattening is reminiscent of the α1 receptor-mediated notion of a ‘network reset’^40^. By increasing response gain (Fig. 1A) through the modulation of second-messenger cascades^4^, noradrenaline released by the LC would augment inter-regional coordination^30^. Importantly, this capacity could confer adaptive benefits across a spectrum, potentially facilitating the formation of flexible coalitions in precise cognitive contexts^46^, while also forcing a broader landscape flattening (i.e., a ‘reset’) in the context of large, unexpected changes^27,40^. Similarly, the idea that phasic cholinergic bursts deepens energy wells is consistent with the idea that the cholinergic system instantiates divisive normalization within the cerebral cortex^14^. Numerous cognitive neuroscience studies have shown that heightened acetylcholine levels correspond to improvements in attentional precision^8,8^. By deepening energy wells, acetylcholine from the BNM could ensure that the brain remains within a particular state and is hence not diluted by other (potentially distracting) brain states. Determining the specific rules that govern the links between the neuromodulation of the energy landscape and cognitive function^47–49^ is of paramount importance, particularly given the highly integrated and degenerate nature of the ascending arousal system^50^.

Our results also provide a systems-level perspective on an emerging corpus of work that details the microscopic circuit level mechanisms responsible for conscious phenomena^51^. In particular, a number of recent studies have highlighted the intersection between the axonal projections of the ascending arousal system and pyramidal cell dendrites in the supragranular regions of the cerebral cortex as a key site for mediating conscious awareness. For instance, optogenetic blockage of the connections between the cell bodies and dendrites of thick-tufted layer V pyramidal cells in the sensory cortex causally modulated conscious arousal in mice^52^. Other work has shown that both the noradrenergic^53^ and cholinergic^54^ systems alter this mechanism, albeit in distinct ways: noradrenaline would promote burst firing due to the α2a receptor-mediated closure of *Ih* HCN leak-channels^53^, whereas the cholinergic system instead prolongs the time-scale of firing via M1 cholinergic receptor activation on pyramidal cell dendrites^54^. In this way, coordinated activity in the ascending arousal system can mediate alterations in microcircuit processing that ultimately manifest as alteration in macroscopic brain network dynamics.

The vascular nature of the T2* fMRI signal is such that it is impossible to rule out the role of haemodynamics in the results we obtained in our analysis. Indeed, there is evidence that noradrenaline causes a targeted hyperaemia through the augmentation of G-protein-coupled receptors on vascular smooth muscle cells^55,56^. However, it is also clear that the haemodynamics and massed neural action in the cerebral cortex are inextricably linked^57,58^. In addition, there is evidence that stimulation of the locus coeruleus leads to the high-frequency, low-amplitude electrophysiological activity patterns characteristic of the awake state^10^. Together, these results argue that the locus coeruleus mediates a combination of haemodynamic and neural responses that facilitate integrative neural network interactions and subsequently mediate alterations in conscious awareness.

In this manuscript, we have argued that the ascending arousal system provides crucial constraints over normal brain function, however there are numerous examples wherein pathology within the ascending arousal system leads to systemic impairments in cognition. In addition to disorders of consciousness^59^, dementia syndromes are also crucially related to dysfunction within the ascending arousal system. For instance, Alzheimer’s disease has been linked to tau pathology within the BNM^26^, however individuals with Alzheimer’s disease also often have pathological involvement of the LC as well^60^. Similarly, individuals with Parkinson’s disease often have extra-dopaminergic pathology in the LC^61^, as well as in the cholinergic tegmentum^62^. Given the pathological processes at play in these disorders, we expect that other neuromodulatory systems will also be impaired, and in turn effect the macroscopic dynamics of the system in ways that remain to be elucidated.

In conclusion, we leveraged a high-resolution 7T resting state fMRI dataset to test the hypothesis that activity within the ascending arousal system shapes and constrains patterns of systems-level network reconfiguration. Our results support specific predictions from a recent hypothetical framework^15^, and further delineate the manner in which the autonomic nervous system shapes and constraints ongoing, low-dimensional brain state dynamics in the central nervous system in a manner that supports changes in conscious awareness.

## Methods

### 7T resting state fMRI

Sixty-five healthy, right-handed adult participants (mean, 23.35 years; SD, 3.6 years; range 18—33 years; 28 females) were recruited, of whom 59 were included in the final analysis (four participants were excluded due to MR scanning issues, one participant was excluded due to an unforeseen brain structure abnormality, and one was excluded due to inconsistent BOLD dynamics following global-signal regression). Participants provided informed written consent to participate in the study. The research was approved by The University of Queensland Human Research Ethics Committee. These data were originally described in Hearne et al., 2017^63^. 1050 (~10 minutes) whole-brain 7T resting state fMRI echo planar images were acquired using a multiband sequence (acceleration factor = 5; 2 mm^3^ voxels; 586 ms TR; 23 ms TE; 40° flip angle; 208 mm FOV; 55 slices). Structural images were also collected to assist functional data pre-processing (MP2RAGE sequence — 0.75 mm^3^ voxels 4,300 ms TR; 3.44 ms TE; 256 slices).

DICOM images were first converted to NIfTI format and realigned. T1 images were reoriented, skull-stripped (FSL BET), and co-registered to the NIfTI functional images using statistical parametric mapping functions. Segmentation and the DARTEL algorithm were used to improve the estimation of non-neural signal in subject space and the spatial normalization. From each grey-matter voxel, the following signals were regressed: linear trends, signals from the six head-motion parameters (three translation, three rotation) and their temporal derivatives, white matter, and CSF (estimated from single-subject masks of white matter and CSF). The aCompCor method (Behzadi et al., 2007) was used to regress out residual signal unrelated to neural activity (i.e., five principal components derived from noise regions- of-interest in which the time series data were unlikely to be modulated by neural activity). Participants with head displacement > 3 mm in > 5% of volumes in any one scan were excluded (*n* = 5). A temporal band pass filter (0.01 < *f* < 0.15 Hz) was applied to the data.

### Brain parcellation

Following pre-processing, the mean time series was extracted from 400 pre-defined cortical parcels using the Schaefer atlas (Schaefer et al., 2018). Probabilistic anatomical atlases were used to define the location of the noradrenergic LC^64^ and the cholinergic BNM (Ch4 cell group)^65^. The mean signal intensity from each region was extracted and then used for subsequent analyses. To ensure that the BOLD data were reflective of neuronal signals, we statistically compared LC and BNM time series with a number of potential nuisance signals from: i) the cerebrospinal fluid; ii) the cortical white matter; iii) mean framewise displacement; and iv) a 2mm^3^ sphere in the fourth ventricle (centred at MNI co-ordinates: 0 −45 −30)^66^. All signals were unrelated to LC and BNM activity (| r | < 0.05 in each case), however given the spatial proximity of the LC to the fourth ventricle, we opted to use a linear regression to residualize the signal from the fourth ventricle. To ensure that BOLD signals from nearby grey matter structures were not influencing the locus coeruleus timcscrics, we extracted the mean activity of the locus coeruleus mask after shifting the mask anteriorly such that it overlapped with an area of the pons that harbours the nuclei (i.e., +8mm in the Y direction). In the same manner in which we previously regressed the dynamics of the fourth ventricle, we regressed the activity of this non-LC pontine region, and then re-analysed our data. Each of the results was statistically identical following this approach, providing confidence that the original conclusions were not biased by a lack of regional specificity.

### Phasic increases in neuromodulatory BOLD signal

To identify phasic increases in neuromodulatory BOLD signal, we calculated the second derivative (i.e., the acceleration) of the LC and BNM time series, and then identified points in time that fulfilled three criteria: 1) value greater than or equal to 2 s.d. above the mean acceleration; 2) value of the original time series, i.e., LC or BNM, was greater than or equal to 2 s.d. above the mean of the time series within the following 10 TRs (i.e., 5.8 seconds); and 3) the time point was not present within the first or last 20 TRs of an individual subjects’ trial (so as to avoid potential boundary effects). Using these criteria, we identified 148 τ_LC-BNM_ time points, 130 τ_BNM-LC_ time points and 316 τ_LC+BNM_ time points across all 59 subjects. To ensure that the choice of 2 s.d. threshold was reflective of the underlying dynamics, we altered this threshold between 1-3 s.d. and found robustly similar patterns. For subsequent analyses, we identified time points in the 21 TR window surrounding these peaks, and then used these to conduct statistical comparisons of the low-dimensional, complex network signature of brain network dynamics as a function of phasic ascending arousal system activity. Each of these patterns was confirmed using a lag-based cross-correlation analysis, which demonstrated similar phenomena to those that we present in the manuscript.

To monitor the propagation of cortical signals with respect to τ_BNM-LC_, τ_BNM-LC_ and τ_LC+BNM_, we extracted the time-to-peak of the cross-correlation between these signals and each of the 400 cortical parcels within the 10 TR (i.e., 5.8 second) windows following each identified phasic peak. These patterns were mapped onto the cortex (Fig. 1B) for visualization and clearly demonstrated anterior-to-posterior direction for the wave. We then used the volumetric MNI co-ordinates of the Schaefer parcellation scheme to calculate the average velocity of the travelling wave (0.13m s^-1^).

In order to obtain an appropriate null model against which to compare our data, we identified 5,000 random timepoints within the concatenated dataset that did not substantially overlap with the already identified τ_LC-BNM_, τ_BNM-LC_ and τ_LC+BNM_ time series, and used these to populate a null distribution^67^. Outcome measures were deemed significant if they were more extreme than the 95^th^ (or 5^th^) percentile of the null distribution. Crucially, this ensured that our data could not be explained by the characteristic spatial and temporal autocorrelation present in BOLD timeseries data.

### Time-resolved functional connectivity

To estimate functional connectivity between the 400 regions of interest, we used the multiplication of temporal derivatives (MTD) technique. Briefly, MTD is computed by calculating the point-wise product of temporal derivative of pair-wise time series. The resultant score is then averaged over a temporal window, w (a window length of 20 TRs was used in this study, though results were consistent for w = 10—50 TRs).

### Modularity Maximization

The Louvain modularity algorithm from the Brain Connectivity Toolbox (BCT^73^) was used on the neural network edge weights to estimate community structure. The Louvain algorithm iteratively maximizes the modularity statistic, *Q*, for different community assignments until the maximum possible score of *Q* has been obtained:

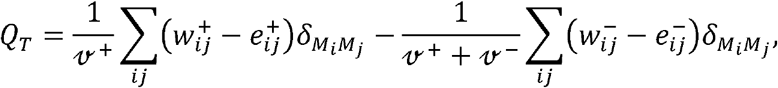

where *v* is the total weight of the network (sum of all negative and positive connections), *w_ij_* is the weighted and signed connection between regions *i* and *j*, *e_ij_* is the strength of a connection divided by the total weight of the network, and *δ_MiMj_* is set to 1 when regions are in the same community and 0 otherwise. ‘+’ and ‘−’ super-scripts denote all positive and negative connections, respectively. The modularity of a given network is therefore a quantification of the extent to which the network may be subdivided into communities with stronger within-module than between-module connections.

For each epoch, we assessed the community assignment for each region 500 times and a consensus partition was identified using a fine-tuning algorithm from the Brain Connectivity Toolbox (BCT; http://www.brain-connectivity-toolbox.net/). We calculated all graph theoretical measures on un-thresholded, weighted and signed connectivity matrices^73^. The stability of the γ parameter was estimated by iteratively calculating the modularity across a range of γ values (0.5-2.5; mean Pearson’s r = 0.859 +-0.01) on the time-averaged connectivity matrix for each subject — across iterations and subjects, a γ value of 1.0 was found to be the least variable, and hence was used for the resultant topological analyses.

### Participation Coefficient

The participation coefficient, *PC*, quantifies the extent to which a region connects across all modules (i.e., between-module strength) and has previously been used to successfully characterize hubs within brain networks (e.g. see^75^). The PC for each region was calculated within each temporal window as,

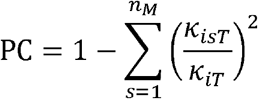

where k_isT_ is the strength of the positive connections of region *i* to regions in module *s* at time *T*, and *k*_iT_. is the sum of strengths of all positive connections of region *i* at time *T*. Negative connections were discarded prior to calculation. The participation coefficient of a region is therefore close to 1 if its connections are uniformly distributed among all the modules and 0 if all of its links are within its own module.

### Brain State Displacement and the Energy Landscape

To quantify the change in BOLD activity following phasic bursts of neuromodulation we calculated the BOLD mean-squared displacement (MSD). The MSD is a measure of the deviation in BOLD activity, ***x**_t_* — [*x*_1,*t*_, *x*_2,*t*_, …, *x_r,t_*] for *r* parcels, with respect to the activity at the phasic onset, *t*_0_. The MSD is calculated as the average change of each voxel

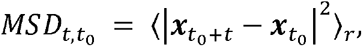

and it is calculated for different ***t*_0_**, where ***t*_0_** are the onset of a subcortical phasic burst, across ***t*** TRs. We are interested in the probability, **P_*MSD*_**, that we will observe a given displacement in BOLD at a given time-lag *t*. We estimated the probability distribution function ***P*(*MSD, t*)** from ***n MSD*_*t, t*_0__** samplings, — e.g., the identified *n* phasic bursts of subcortical structures (as above) — using a Gaussian kernel density estimation 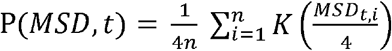 where 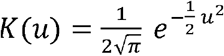 and we display the results for ***t*** between 1 to 15 TR and ***MSD*** between 0 to 50. As is typical in statistical mechanics the energy of a given state, ***E_σ_***, and its probability are related 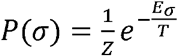, where ***Z*** is the normalisation function and ***T*** is a scaling factor equivalent to temperature in thermodynamics^20^. In our analysis **∑_*σ*_ *P_σ_* = 1** by construction and we can set ***T* = 1** for the observed data. Thus, the energy of each BOLD MSD for a given at a given time-lag ***t*, E**, is then equal to the natural logarithm of the inverse probability, ***P***(***MSD, t***), of its occurrence:

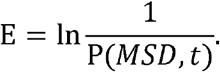

### Meditation Dataset

Fourteen healthy right-handed non-smoking meditation practitioners (11 female; age 28-66) underwent Siemens 3T MRI scanning (T1: TR = 2600 msec, TE = 3.9 msec, TI = 900 msec, FOV = 24 cm, 256 x 256 matrix, voxel dimensions =1×1×1 mm^3^; 12*: weighted gradient-echo pulse sequence, TR = 1500 msec, TE = 30 msec, flip angle = 90 deg, FOV =192 cm, 64 x 64 matrix, voxel dimensions = 3×3×4 mm^3^). All participants signed a consent form approved by the Institutional Review Board at Emory University and the Atlanta Veterans Affairs Research and Development Committee as an indication of informed consent. Participants were asked to meditate for 20 min in the MRI scanner by maintaining focused attention on the breath and keeping the eyes closed. They were instructed to press a button whenever they realized their mind had wandered away from the breath, and then return their focus to the breath. The epoch of time immediately prior to the button press was thus the moment in time in which each individual recognized that their focus had deviated from their breath. This information was used to construct a finite impulses response model that mapped the 5 TRs prior-to and following each button press. We then modelled LC>BNM activity, low-dimensional dynamics and network topology around this epoch to construct a model of state-space reconfiguration as a function of intrinsic conscious awareness. Non-parametric, block-resampling null distributions were utilized for statistical testing (p < 0.05).

## Data availability

The BOLD data was obtained from (Hearne et al., 2017)^63^ and The BOLD data that support the findings of this study were obtained from (Hearne et al., 2017)^63^ and they are available from the authors upon reasonable request. The subcortical tìmcscrics (*τ_LC_* and *τ_BNM_*) that support the findings of this study are available at (github.com/Bmunn/BSI).

## Code availability

All the code required to conduct the analysis can be found on Github at (github.com/Bmunn/BSI).

## Supplementary Figures

**Figure S1.**
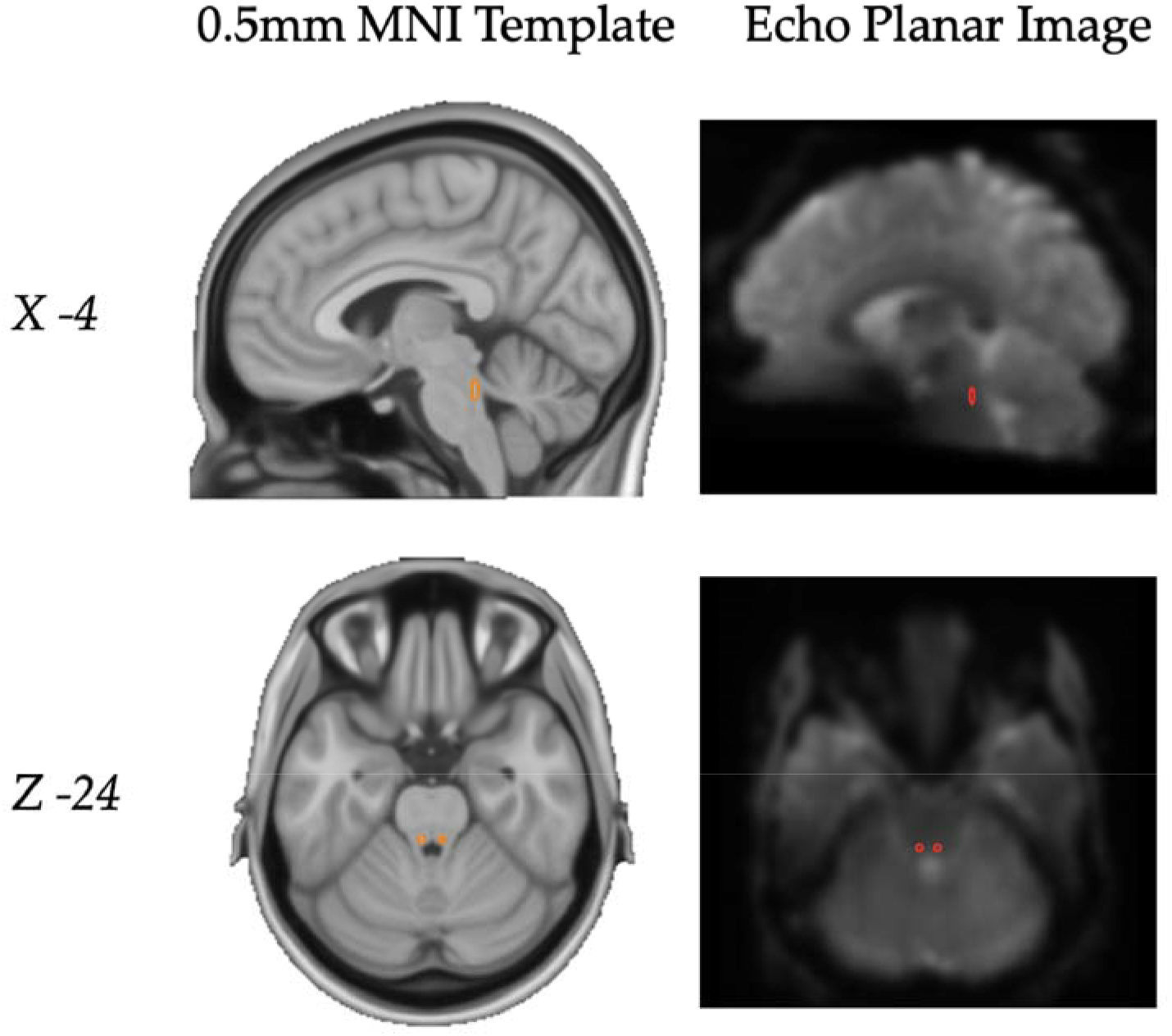
The locus coeruleus. Left: The anatomical locus coeruleus mask projected onto MNI 0.5mm standard brain (orange); Right: The anatomical locus coeruleus mask down-sampled onto an example 7T Echo Planar Image from a single subject (red).

**Figure S2.**
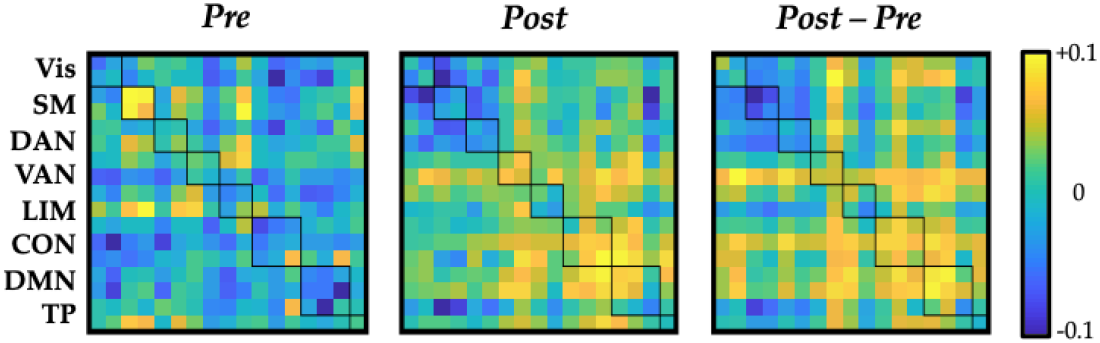
Time-varying correlations. Average correlation preceding (left) and following (middle) the zero- lagged value, along with the difference between the two (right); squares represent eight pre-defined sub-networks: Vis — visual, SM — somatomotor, DAN — dorsal attention, VAN — ventral attention, L1M — limbic, CON — control, DMN — default and TP — temporal pole.

**Figure S3.**
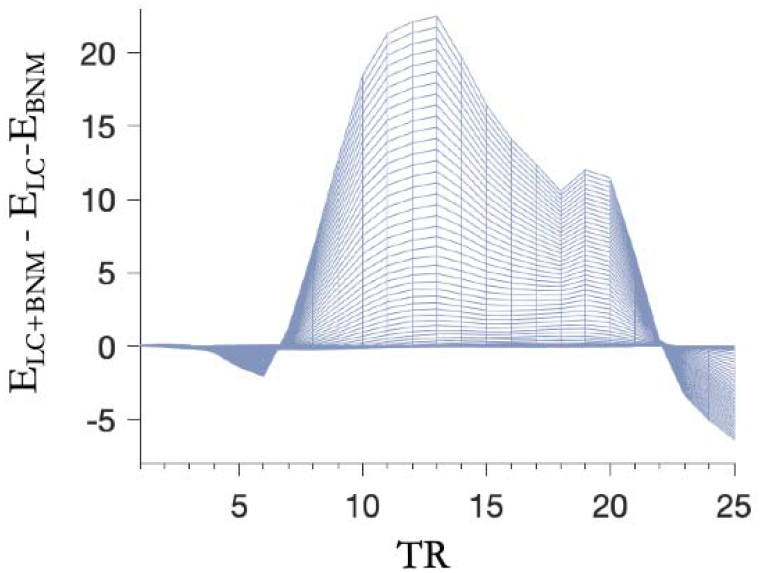
LC and BNM move dynamics to differing regimes than unaroused activity and their simultaneous combination LC+BNM. The energy landscape of simultaneous LC+BNM phasic bursts relative to their linear superposition, suggesting the simultaneous combination may allow the system to reach particularly unique brain-states that neither individually could reach.

## Author contributions

MS conceived, funded, and directed the project. MS Curated the data. BM, EM, MS Conducted the analysis. BM, MS Wrote the original draft. BM, EM, GW, MS Reviewed and edited the manuscript.

